# Obstacle bypassing and intrinsically asymmetric loop extrusion in the segment-capture model of Structural Maintenance of Chromosomes complexes

**DOI:** 10.64898/2026.07.07.736920

**Authors:** Lucas Dooms, Stefanos Nomidis, Enrico Carlon, Stephan Gruber, John F. Marko

**Affiliations:** Soft Matter and Biophysics, KU Leuven, Celestijnenlaan 200D, B-3001, Leuven, Belgium; Département de Microbiologie Fondamentale, Université de Lausanne, 1015 Lausanne, Switzerland; Department of Physics and Astronomy, and Department of Molecular Biosciences, Northwestern University, Evanston, Illinois 60208, USA

## Abstract

Structural Maintenance of Chromosomes (SMC) complexes are large protein assemblies that play a central role in chromosomal organization across all domains of life by acting as molecular motors that extrude DNA loops. Although the molecular architecture of SMCs has been characterized in considerable detail, their mechanistic mode of action remains actively debated. One proposed mechanism is the segment-capture model, in which SMC complexes undergo an asymmetric conformational change upon ATP binding and hydrolysis, effectively “pumping” short DNA segments through the SMC ring and thereby enlarging loops. While this model accounts for several experimentally observed features of loop extrusion, it has been questioned in light of experiments showing that SMC complexes can traverse large nanoparticle-sized obstacles on DNA. Moreover, recent experimental advances have provided new insights that extend beyond the original formulation of the model. Here we show that, contrary to previous criticisms, the segment-capture model can naturally account for key features of recent experiments, including the bypass of large obstacles when the SMC hinge acts as a bypass gate. Furthermore, we outline a set of targeted experiments designed to critically test the model’s predictions, providing a clear path toward resolving outstanding questions about the mechanistic basis of loop extrusion.

## 1 Introduction

Structural Maintenance of Chromosomes (SMC) complexes constitute a class of protein complexes that are essential for the compaction and higher-order organization of chromosomes in both eukaryotes and bacteria. In eukaryotic cells, the principal SMC complexes are cohesin, condensin, and SMC5/6, which play central roles in chromosome compaction, segregation, and DNA repair. Bacteria likewise possess SMC complexes, such as bsSMC in *Bacillus subtilis* and the MukBEF complex in *Escherichia coli*, and use the SMC-based defense system Wadjet for plasmid restriction.

All SMC complexes display a high degree of structural similarity and are strongly conserved across organisms from both the eukaryotic and prokaryotic domains of life. In SMC complexes, the protein subunits assemble into a tripartite ring (Fig. 1) comprising two long coiled-coil arms joined at a hinge region, with ATP-binding “head” domains located at the opposite ends. These head domains are bridged by a kleisin subunit. The kleisin is relatively short in prokaryotic SMC complexes and in the eukaryotic SMC5/6 complex (approximately 250 amino acids), but is substantially longer due to a large flexible center in eukaryotic cohesin and condensin (on the order of 500 amino acids). Various regulatory subunits associate with the kleisin, and multiple experimental studies have demonstrated that these complexes are capable of binding to and topologically entrapping DNA strands [1– This body of evidence supports earlier models proposing that DNA is topologically enclosed within the cohesin SMC ring, thereby maintaining the association of sister chromatids [6] (throughout this paper, the term “strand” will refer to a DNA duplex or dsDNA, except when the term “singlestranded DNA” or ssDNA is used, which refers to one polynucleotide chain).

**Figure 1:**
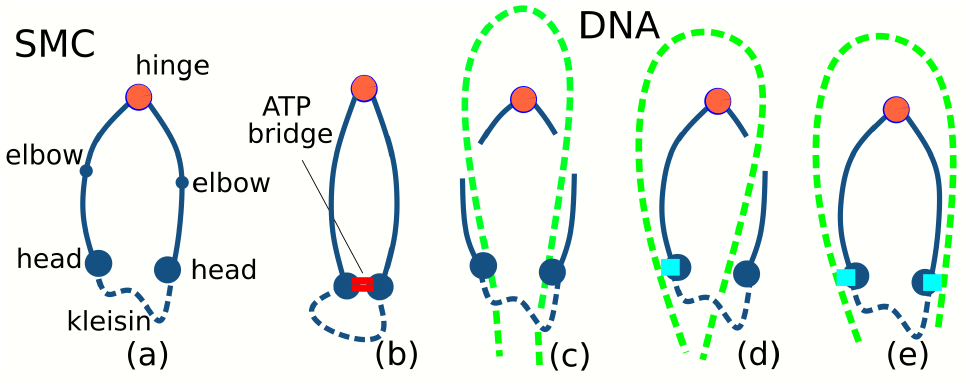
(a) The common basic architecture of Structural Maintenance of Chromosomes (SMC) protein complexes is that of a tripartite ring. This is formed by two coiled-coil proteins (blue, solid) which are joined to each other at a hinge (orange) and bound to a kleisin protein (blue, dashed) at the head sites. (b) Upon ATP binding the two heads form a bridge (red) which subdivides the SMC ring into two compartments. (c-e) Alternative modes of interaction between SMC and a DNA molecule (green, dashed) to produce loop extrusion. These are referred to as (c) pseudo-topological, (d) topological and (e) non-topological, with two, one or no DNA strands threaded through the SMC ring. This paper discusses the segment-capture model, which can describe both pseudotopological (c) and topological (d) loop extrusion.

The role of complexes like cohesin and condensin in chromosome organization and dynamics is well established. In vitro experiments have shown that these complexes act as molecular motors, translocating along DNA [7] and extruding loops [8]. In working as molecular motors, SMCs cycle through different conformations via ATP binding and hydrolysis. The head domains contain ATP binding sites and when two ATP molecules are bound a bridge is formed and the SMC ring is partitioned in two compartments (Fig. 1(b)). These conformational changes should be linked to loop extrusion; however, precisely how this occurs is a topic of current debate, and different models have been proposed.

As illustrated in Fig. 1(c-e), loop extrusion (LE) by SMC complexes can be categorized into three principal mechanistic classes: (c) pseudo-topological, (d) topological, and (e) non-topological. In the pseudo-topological scenario (c), both arms (sides) of the DNA loop are threaded through the SMC ring. In contrast, in the topological case (d), only one arm of the DNA loop is inserted into the SMC ring lumen, while the other remains bound to the exterior surface of the SMC complex or, in the case of dimeric SMC motors such as Wadjet and MukBEF, is inserted into the lumen of the mate pair SMC complex [3, 5, 9]. The third mechanistic class is non-topological LE (e), in which both DNA arms interact with the complex from the outside and are not topologically trapped within the ring. Earlier theoretical and experimental studies predominantly considered topological or pseudo-topological LE, i.e., mechanisms in which at least one of the two DNA strands at the base of the loop is threaded through the SMC ring. More recently, however, non-topological LE has been proposed to account for single-molecule in vitro observations [10].

To emulate LE in chromatin within living cells, where DNA is extensively associated with proteins and large macromolecular complexes, these experiments systematically probed the influence of various DNA-bound “obstacles”. It was found that, while relatively small factors such as nucleosomes or RNA polymerases can plausibly pass through the SMC lumen, LE persisted even in the presence of DNA-bound obstacles whose size substantially exceeds the ring diameter [10]. On this basis, the authors inferred that a non-topological mechanism is the only viable explanation for the observed robustness of LE. This conclusion, however, has been challenged by a study demonstrating that the SMC hinge can act as a gate that enables bypass of obstacles that are tethered to double-stranded DNA (dsDNA) via single-stranded RNA (ssRNA) or ssDNA linkers [11].

While the microscopic details through which such a non-topological LE would work are not yet clear, some models have been presented [12]. These suggest that the SMC walks along a DNA molecule without topologically embracing it inside the ring, possibly folding the peptide chain of the kleisin complex around the DNA [12, 13]; see the Discussion for more details. In Fig. 1, DNA binding sites for DNA outside the lumen are indicated in light blue in the topological (d) and non-topological (e) model. We note that these have different functions in the two cases. While in the topological model the binding site can remain a passive DNA anchoring site, at least one of the two sites in the non-topological case must “walk along the DNA” in order to produce LE. We note that the exact positions of the two binding sites in the non-topological model (Fig. 1) are not known. If these sites are expected to exhibit motor activity, we assume they are located in close proximity to the ATP-binding sites, near the SMC heads or the kleisin. Although the non-topological model is appealing in that it can account for the circumvention of large obstacles, it presents several challenges, which will be examined in greater detail in subsequent sections of this paper.

### 1.1 (Pseudo) topological segment-capture model

A topological (or pseudo-topological) mechanism for LE is provided by the so-called segment-capture model [14, 16, 17]. The molecular details of the LE mechanism within this framework have been elucidated, and a coarsegrained molecular dynamics (MD) simulation code has been developed [14] (Fig. 2).

**Figure 2:**
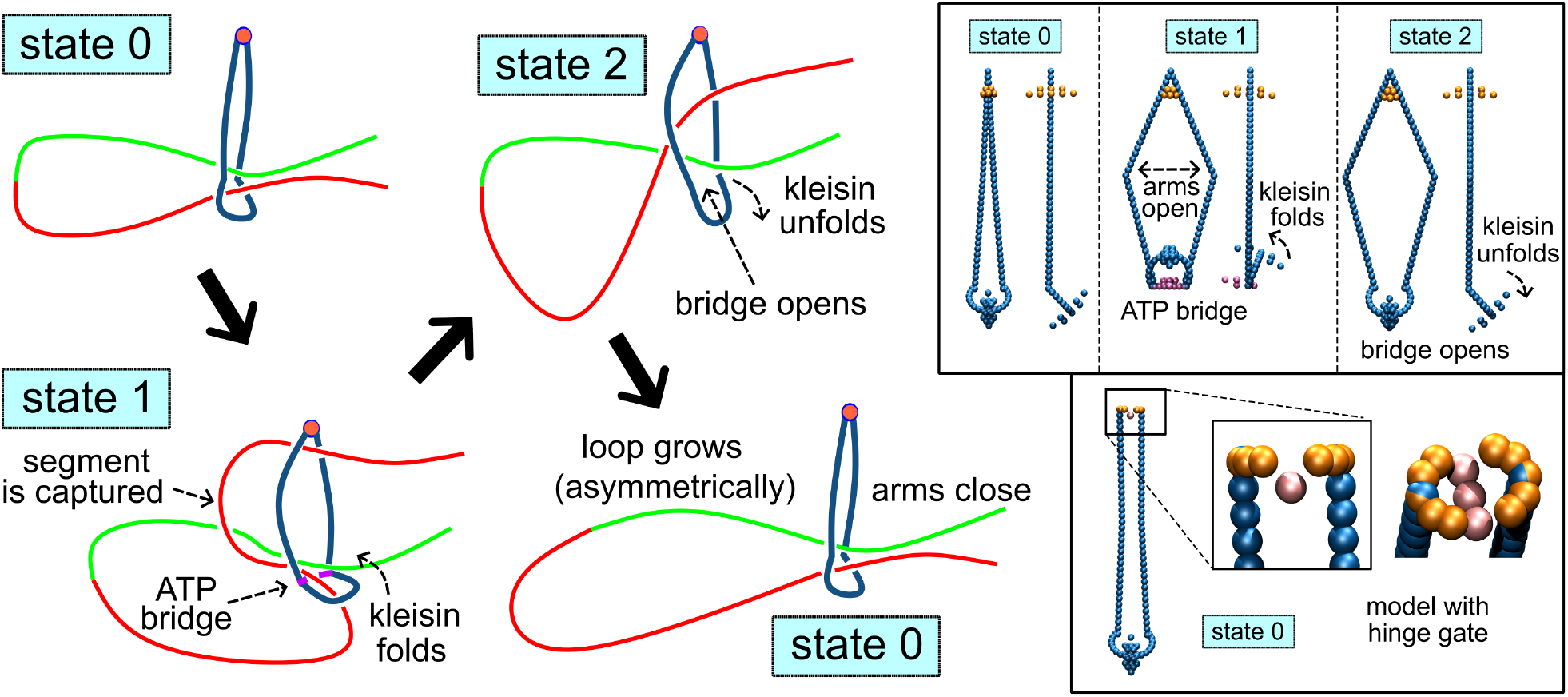
In the segment-capture model of SMC loop extrusion dynamics, the SMC ring (in blue) undergoes conformational changes cycling through three states 0 *→* 1 *→* 2 *→* 0 *→* 1 *→ · · ·*. Here we show a pseudo-topological loading of the DNA into the SMC (state 0, top left). In this state the SMC arms are closed. In state 1 an ATP bridge is formed and the SMC arms are open. A strand is trapped in the kleisin, which upon ATP binding folds asymmetrically and pushes a segment of DNA through the SMC lumen. In state 2 the bridge opens and the captured segment is released, contributing to the growth of the loop. The SMC returns to state 0 and the DNA is pushed back towards the kleisin. For clarity we have divided the DNA into two halves of different colors. In the example the red part is captured and contributes to the growth of the loop, which grows asymmetrically. A mechanism by which directional switching produces symmetric loop extrusion is discussed in the paper. The top right frame shows the actual MD model with details of the three states, following the model of Ref. [14]. The bottom right frame shows a version of the SMC containing a hinge gate, used to study bypass of large obstacles. MD snapshot images were produced using VMD [15].

In our MD implementation of the segment-capture model, the SMC complex undergoes cyclic transitions between three distinct conformational states in which the main ring lumen is either open or closed, and the kleisin subunit can adopt two distinct folded conformations. At the beginning of the cycle, the DNA is topologically entrapped within the kleisin compartment, while the SMC ring is in a closed conformation with the two coiled-coil arms in close proximity. Upon ATP binding, a bridge is formed and the kleisin folds, thereby forcing a short DNA segment to enter the SMC ring (hence the designation “segment-capture”). Note that the captured segment forms a small arc of DNA (in itself a small loop) on the order of a few hundred bp. Following ATP hydrolysis (ATP *→* ADP), the bridge is removed, and the portion of the captured DNA segment that was previously confined within the kleisin compartment is released. This contributes to the growth of the loop. At the end of the cycle, the two coiled-coil arms re-approach each other, displacing the DNA back toward the kleisin compartment and thereby restoring the initial configuration. In this model, the DNA remains topologically entrapped within the SMC ring throughout the cycle.

Coarse-grained molecular dynamics (MD) investigations of this mechanism have been reported in Refs. [14, 18], providing insights into several key features of the LE process at different levels of coarse graining. In particular, the model of Ref. [18] employs a finer level of coarse graining than that of Ref. [14], enabling a more detailed molecular characterization of DNA-SMC interactions throughout the different stages of the LE cycle. In contrast, the coarser model of Ref. [14] is better suited to follow loop extrusion over longer time scales and across a larger ensemble of trajectories, thereby yielding robust statistics and a more accurate characterization of the stochastic properties of the LE process.

### 1.2 Aim of the paper

In recent years, a number of experiments have provided new insight into the LE activity of SMC complexes, extending well beyond what was known at the time the segment-capture model was first proposed and developed [14, 16, 17]. This growing body of experimental evidence motivates further analysis of the segment-capture model. The newly available data invite us to test the assumptions and predictions of the model, in light of direct molecular observations. The aim of this paper is therefore to focus specifically on the segment-capture model, to reevaluate its validity and limitations given the most recent singlemolecule experimental results, and to discuss alternative mechanisms for non-topological LE models.

## 2 MATERIALS AND METHODS

### 2.1 Molecular dynamics model

The coarse-grained implementation of the segment-capture model employed in this study is essentially identical to that described in Ref. [14], with the additional inclusion of a hinge gate that was absent in the original formulation. The SMC complex is represented as an assembly of rigid bodies that interact via angular and dihedral potentials. In this coarse-grained framework, the SMC cycles through three distinct conformational states, denoted “0”, “1”, and “2”, as depicted in the top right panel of Fig. 2. These states correspond to the ATP-unbound (apo), ATP-bound, and ATP-hydrolyzed/ADP-bound states of the ATPase, respectively. The transitions 0 *→* 1 *→* 2 *→* 0 *→* … are implemented as instantaneous changes in the potential energy function, such that the equilibrium angles between the rigid elements are abruptly modified. Transitions from state *i* to state *j* (with *i, j* = 0, 1, 2) occur with rates *k*_*i→j*_, with only *k*_0*→*1_, *k*_1*→*2_ and *k*_2*→*0_ non-vanishing. In some simulations we enforced transitions at fixed time intervals Δ*t*_*i→j*_ = 1*/k*_*i→j*_. Within each state, the system evolves according to standard molecular dynamics (MD) with an implicit-solvent Langevin thermostat implemented in LAMMPS [19]. Further methodological details, as well as the structural evidence underpinning this model, are provided in Ref. [14]. The specific adaptations introduced in the present work are summarized briefly below.

### 2.2 Hinge gate

A recent study [11] demonstrated that the hinge functions as a selective bypass gate capable of accommodating single-stranded nucleic acid linkers, such as ssDNA, but not double-stranded DNA. To incorporate this mechanism, we developed a modified version of the MD segment-capture model that includes a bypass gate at the hinge, i.e., at the junction of the two coiled-coil arms of the SMC complex. In the coarse-grained model introduced in Ref. [14], the hinge is represented by two rigid segments connected via a single shared coarse-grained bead. In the new implementation, by contrast, the hinge region is modeled as two rigid semicircles maintained at a fixed separation.

The two panels on the right-hand side of Fig. 2 depict these alternative hinge representations: the original model (top), which includes all three conformational states, and the modified model with a hinge gate (bottom), for which only state “0” is illustrated. The gap width between the two semicircles is chosen such that the larger effective diameter of dsDNA prevents it from traversing the gate, whereas the smaller diameter of ssDNA allows it to pass. We examined the impact of varying the gap width by considering two distinct distances between the semicircles.

### 2.3 DNA, obstacles and nucleosomal particles

DNA was modeled, following Ref. [14], as a semiflexible polymer using a coarse-grained representation in which each bead corresponds to 5 bp and interacts via excluded-volume repulsion. Adjacent beads are connected by Finite Extensible Nonlinear Elastic (FENE) bonds, and consecutive bonds are subject to a bending energy of the form *E* = *K* (1 + cos *θ*), where *θ* denotes the bending angle. The parameter *K* is a stiffness coefficient that we systematically varied in order to modulate the effective DNA persistence length within the SMC translocation process. Obstacles were modeled as single beads interacting through a soft repulsive WCA potential. To reproduce the conditions of the experiments, obstacles were assigned different sizes and modes of attachment to the DNA. Obstacles with diameters in the range 10 nm *≤D ≤*22 nm were directly incorporated into the DNA polymer to emulate nucleosomes, RNA polymerase, or dCas9 used in experiments [10]. A larger obstacle with diameter *D* = 200 nm was attached to the DNA via a linker molecule, again following the experimental setup [10, 11]. This tether consisted of an ssDNA segment modeled as a modified ds-DNA strand with half the diameter, a reduced persistence length of 1 nm, and no interactions with the SMC binding sites, while all other dsDNA properties were kept unchanged for simplicity.

## 3 RESULTS

We present the results of molecular dynamics (MD) simulations conducted within the framework of the segmentcapture model.

### 3.1 Directional switching

It has been established that some SMC complexes extrude DNA loops in an asymmetric manner, while others operate symmetrically. Single-molecule experiments [8, 20] have demonstrated that condensin extrudes loops with one DNA segment remaining anchored while the other is reeled in by the motor activity of the complex, yielding loops that grow from only one side. Condensin is therefore classified as an asymmetric loop-extruding factor. Recent work showed that deletion of specific HEAT-repeat proteins associated with the kleisin subunit converts condensin into a symmetric loop extruder [21]. Initial in vitro single-molecule assays indicated that cohesin extrudes loops symmetrically [20, 22, 23]. However, more recent single-molecule experiments indicate that cohesin and SMC5/6 primarily extrude DNA loops asymmetrically. These complexes frequently undergo strand switching during LE, so that over sufficiently long observation times their activity may appear effectively symmetric [24].

We show here that the segment-capture model can incorporate directional switching in a simple and natural manner, using a mechanism proposed in Ref. [14]. For this purpose, we use a pseudo-topological LE mechanism in combination with internal strand-swap events, see Fig. 3. To elucidate the mechanism, we represent the extruded DNA loop with two differently colored halves, red and green, with the green strand bound to the inner site of the SMC lumen (Fig. 3(a)). If the green segment were to remain permanently bound, the SMC complex would cycle through the conformational states 0 *→* 1 *→* 2 *→* 0 while acting exclusively on the red strand (see Fig. 2), thereby generating an asymmetric LE process in which the loop grows from the red strand side [see SI movie S1]. Transient unbinding of the green strand can give rise to a strand-swapping event (Fig. 3(c)), in which the red strand becomes bound instead. When the red strand is bound, the SMC acts on the green strand, and the loop grows from the green strand side.

**Figure 3:**
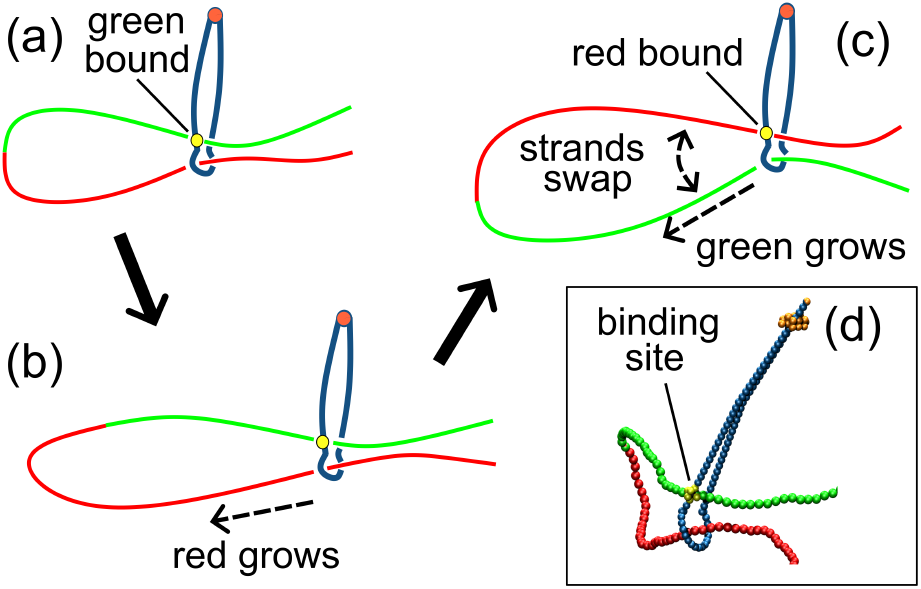
(a-c) Schematic mechanism of strand switching in LE. In the model only one of the two strands is trapped in the kleisin compartment, while the other is kept outside the kleisin compartment and bound to a DNA binding site inside the SMC ring (a). If the strand remains permanently bound to the SMC the LE proceeds asymmetrically (b). A weaker binding site can lead to unbinding and strand swapping (c), leading to a directional switch. Thus every single motor cycle is asymmetric, but a periodic strand swapping leads to a symmetric extrusion. An actual snapshot of the MD simulation is shown in (d).

Frequent strand swapping of this sort leads to a net symmetric extrusion when averaged over many cycles, despite each individual cycle being intrinsically asymmetric [see SI movie S2]. This behavior is in accord with recent single-molecule observations [24]. In the described mechanism both strands are inside the SMC lumen, corresponding to a pseudo-topological model, Fig. 1(c). The case of one strand permanently bound externally to the SMC (Fig. 1(d)) has the same single-cycle dynamics, but cannot switch strands, and produces asymmetric LE as was discussed in [14].

Figure 4 shows a plot of an MD simulation run monitoring the positions *x*_*r*_, *x*_*g*_ of the two DNA strands with respect to the SMC over time. These positions are those of the DNA coarse-grained beads crossing the SMC lumen. In the case of multiple crossings, such as for the red strand in Fig. 2 (state 1), the recorded position *x*_*r*_ is that of the bead closest to the kleisin. The top main panel of Fig. 4 shows a plot of an MD simulation run for a permanently bound green strand. Here only the “red” strand grows (red trace) while the “green” strand remains permanently anchored *x*_*g*_ = const., which leads to a net asymmetric LE. In Fig. 4 the purple dashed trace shows the total size of the loop defined as |*x*_*g*_ *− x*_*r*_|. The vertical dashed lines indicate the switching between the three states of the SMC (0 *→* 1 *→* 2). For the sake of clarity in this run the duration of each state is constant, while in most simulations they are selected randomly from an exponential distribution [14]. According to the original model, state 1, i.e. when the segment is captured (Fig. 2), has the longest average duration. The top inset shows a zoom in of the *x*_*r*_ trace. At the beginning of state 0, while the SMC is closing its arms, the *x*_*r*_ changes slightly and it remains rather constant. In state 1 the DNA is tightly bound to the folded kleisin and *x*_*r*_ does not vary. It is in the state 2 that *x*_*r*_ shows some variations: it increases in steps of *∼*50 nm on average, but it is also characterized by some backward steps [14].

**Figure 4:**
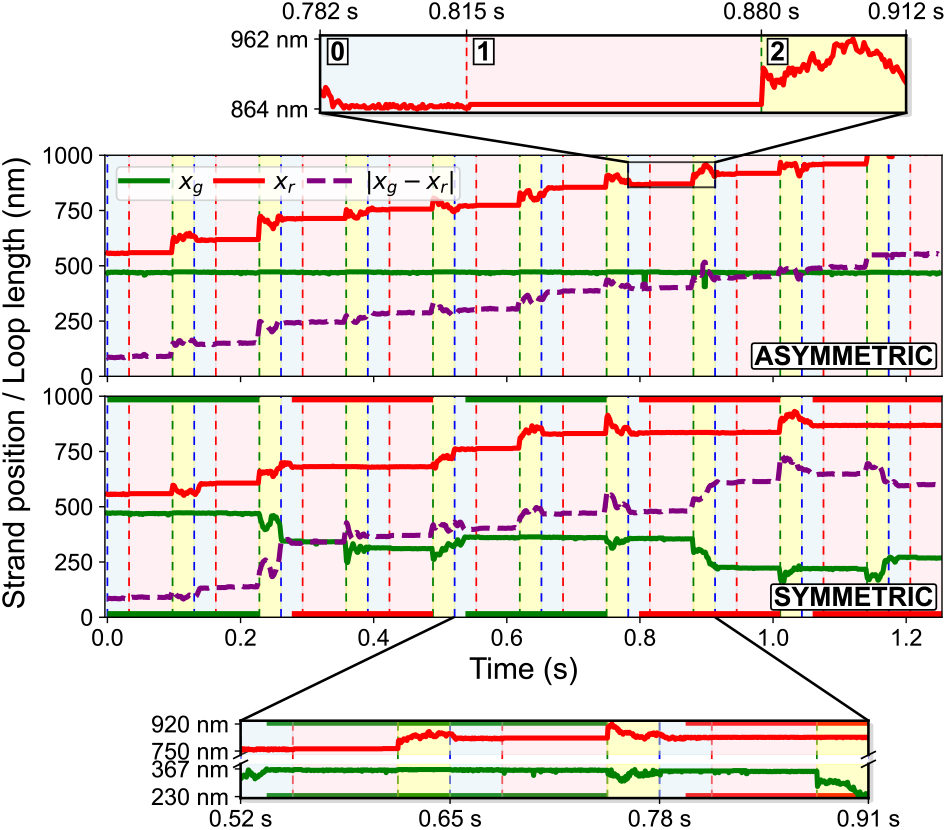
MD traces showing the time evolution of *x*_*g*_ and *x*_*r*_ (positions of the green and red strands as defined in the text) and of the loop length | *x*_*g*_ *− x*_*r*_ |. In the top panel the green strand is permanently bound to the SMC, resulting in fully asymmetric LE. In the bottom panel the bound strand is forced to detach every two SMC cycles, producing a symmetric LE composed of intrinsically asymmetric steps. The color of the bound strand is shown on the *x*-axis. The vertical stripes identify the 0, 1 and 2 SMC states, here taken to have constant durations across cycles (the state 1 is the longest of the three according to the parametrization of Ref. [14]). Two zoom-ins of the top and bottom graphs are shown. Note that the strands grow primarily in state 2.

Figure 4 (bottom main panel) shows an MD simulation run in which we have chosen to enforce the unbinding of the bound strand at the end of SMC state 1 every two cycles. After the bound strand is released and the SMC enters state 0 in the following cycle, a strand of either color can bind to the binding site. The other (nonbound) strand is then trapped in the kleisin and contributes to the loop growth. The specific run shown at the bottom of Fig. 4 shows three strand-swap events, at times *t ≈*0.25s, *t ≈*0.5s, and *t ≈*0.8s, as well as a detachment-reattachment of the same red strand at *t ≈*1.0s. The color of the bound strand is shown as a thick horizontal line along the time axis in the figure. The net result of the strand swapping dynamics is a symmetric extrusion. We note that for short time intervals both strands are unbound from the SMC, which then performs unbiased diffusion along the DNA loop. This loop sliding was observed in a recent single-molecule experiment [24]. Movies of two simulation runs for asymmetric LE [Movie S1] and symmetric LE with strand swap mechanism [Movie S2] are provided in the SI.

In our MD model we constrained one of the two DNA strands at a designated binding site external to the kleisin. The kleisin itself was represented as a rigid semicircular element that undergoes asymmetric folding to generate the motor power stroke [14]. In reality, however, the kleisin contains extended intrinsically disordered or flexible regions that serve as binding platforms for multiple regulatory proteins (see also Discussion). It is therefore more plausible that these regulatory factors mediate the entrapment of one DNA strand (the red strand) and drive the associated folding/power-stroke event, while other flexible kleisin regions could accommodate the other (green) strand. Under this scenario, a dedicated DNA binding site is not required, and the other DNA strand (the green strand) would remain topologically constrained within a portion of the kleisin that does not undergo folding.

### 3.2 Obstacle crossing via linker through the SMC hinge

Loop extrusion in living cells must operate in a highly crowded environment, as chromosomal DNA is bound to a large number of proteins of diverse sizes. Understanding how DNA LE proceeds under these conditions has become a central question, prompting extensive experimental and theoretical investigations [13]. The majority of

DNA-bound proteins in both eukaryotic and bacterial cells have dimensions ranging from a few nanometers (e.g., the histone octamer) to approximately 15–20 nm (e.g., DNA polymerase). In principle, the SMC ring is sufficiently wide to accommodate proteins of this scale in its lumen. Furthermore, the kleisin subunit is substantially longer in cohesin and condensin (*≈* 500 aa) but shorter in SMC5/6 and in some prokaryotic SMC complexes [25]. These differences suggest that, during evolution, SMC complexes may have acquired wider ring architectures to facilitate the passage of larger DNA-bound proteins.

Pradhan et al. [10] conducted a series of singlemolecule experiments to investigate DNA translocation in the presence of various DNA-bound protein complexes, demonstrating that SMC complexes can translocate past nucleosomes, RNA polymerase, and dCas9. A particularly surprising observation was that LE by condensin and cohesin was not stopped in a significant fraction of cases when DNA was tethered to nanoparticles with diameters of up to 200 nm, which substantially exceed the size of the SMC ring (see Fig. 5(a)). These results were initially interpreted as strong support for a non-topological LE model [10], i.e. with DNA not trapped inside the SMC ring, as shown in Fig. 1(e).

**Figure 5:**
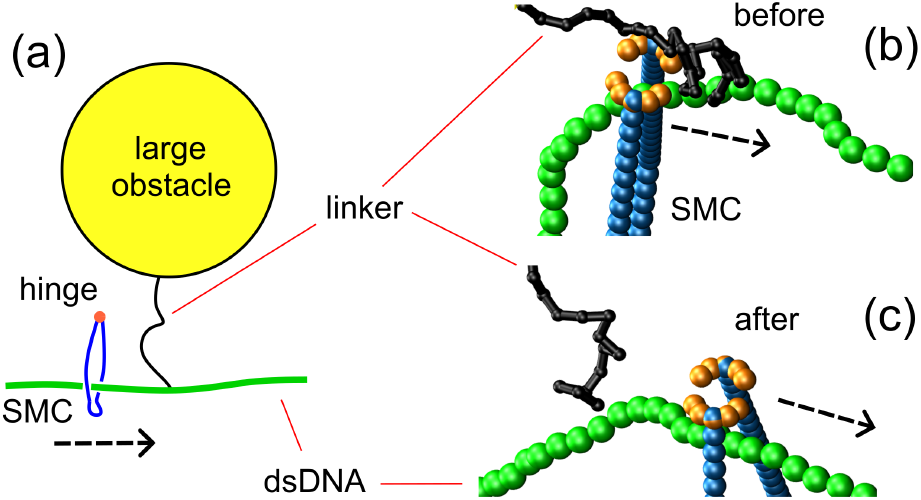
(a) In vitro single-molecule experiments have shown that SMC complexes can traverse large obstacles (such as 200 nm gold nanoparticles), which are much larger than the SMC ring [10]. Evidence from experiments on bacterial SMC complexes suggested that the hinge site (orange) can act as a selective gate that allows the passage of thinner ssDNA or ssRNA, while preventing passage of thicker dsDNA. We extended the MD model of the SMC by adding such a selective gate, modeled as two neighboring rigid semicircles (yellow) sufficiently separated to allow the passage of thin linker polymers, such as ssDNA or ssRNA, but not dsDNA. (b,c) Zoom-ins of an MD simulation event showing the passage of the thin flexible linker (purple) through the hinge gate. The dashed arrows indicate the direction of the translocation of the SMC. The snapshots show configurations (b) just before and (c) immediately after one passage event. To aid visualization, the large 200 nm particle tethered to the linker is not shown in (b) and (c).

However, more recent experiments have proposed an alternative mechanism for obstacle crossing [11]. A crucial aspect is that the obstacle (nanoparticle) is connected to the DNA via a flexible linker molecule (see Fig. 5(a)).

It was argued that bypass occurs because this linker can be selectively captured within the SMC hinge channel, which functions as a gated conduit: it permits thinner linkers, such as ssDNA and ssRNA, to pass through, while constraining and retaining dsDNA within the SMC ring [11]. Consistent with this mechanism, the experiments further demonstrated that obstacle bypass is abolished when thicker dsDNA linkers are employed [11]. We note that the experiments of Ref. [10] involve LE, but for simplicity in our simulations we considered the translocation of a single dsDNA, as in Fig. 5(a). To analyze the properties of obstacles tethered to dsDNA via linker molecules we used the version of the coarse-grained SMC model with a hinge gate, shown in Fig. 2, bottom right panel.

Figure 6 shows 50 individual time traces of the SMC position along the dsDNA for several translocation events under three different conditions, with the thick purple line indicating the average position over all runs, and the gray area representing an interval of *±σ*, with *σ* the standard deviation. Figure 6(a) shows, as a reference, the case of translocation along naked DNA with no obstacles attached. Figures 6(b-d) depict three cases of translocation with obstacles and (b) a wider gate at the hinge site, (c) a narrower gate and (d) a sealed gate. Once the SMC reaches the end of the DNA (position 0 nm) and detaches, the trajectory stops. To compute average velocities beyond this point, we artificially extend the trace according to the average velocity of the naked DNA translocation of Fig. 6(a). The average translocation velocity is *∼*700 nm/s or *∼* 2 kb/s. The linker strand is attached to position 510 nm along the dsDNA, which is shown as a horizontal dashed-dotted line. In the simulations shown in Fig. 6(b), the translocation is almost as fast as that without any obstacle, case (a). In both cases the average SMC position appears to be almost linear in time showing that the average SMC velocity is approximately constant. With a narrower gate (c), the average velocity decreases when the SMC approaches the linker attachment position. The individual trajectories show that many trajectories “bounce” from the obstacle much more frequently than in case (b). Once a trajectory bypasses the obstacle, the hinge gate prevents the SMC from sliding backwards, effectively speeding up the translocation. We note that in the narrow gate simulations of (c) a significant number of trajectories do not cross the obstacle. We estimate that this happens for about 15% of cases. Experimental data indicate that about 50% of 200 nm obstacles are blocked in experiments with cohesin and condensin [10].

**Figure 6:**
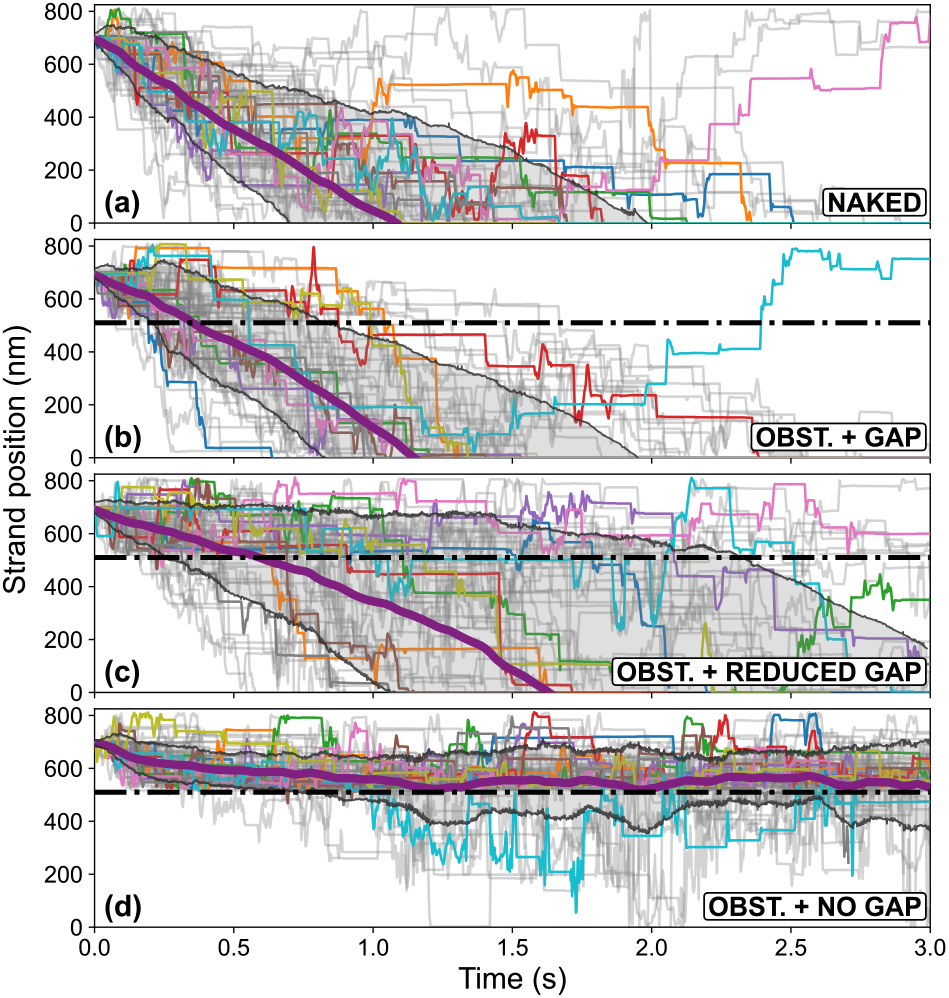
Comparison of translocation along (a) naked DNA and (b-d) DNA with an obstacle tethered via a linker molecule, as shown in Fig. 5(a). The linker molecule is attached to the dsDNA at position 510 nm, shown as a horizontal dashed-dotted line in (b-d). These three cases show (b) a large hinge gap, (c) a small hinge gap, and (d) a closed hinge. In each simulation, the SMC starts at 697 nm and moves towards the end of the dsDNA located at 0 nm. The thin gray lines represent trajectories of individual simulation runs, with 10 randomly chosen trajectories highlighted in color. The thick purple line shows the average SMC position over the individual trajectories and the gray area represents an interval of *±*1 standard deviation.

Complete closure of the hinge gate does not entirely stop translocation: a small fraction of trajectories manage to bypass the obstacle (Figure 6(d)). Figure 7 provides a schematic illustration of typical configurations observed in these cases. When the hinge gate is closed, the linker strand cannot pass through it. In most cases this halts translocation (Fig. 7(a)). However, in some cases the blockage is bypassed (Fig. 7(b)): in these cases, the SMC complex captures and translocates DNA beyond the position of the linker molecule, generating an additional loop. Such behavior was previously hypothesized in Ref. [14]; the present simulations confirm that this represents a feasible mechanism for progression of translocation in the presence of obstacles, even when the hinge gate is sealed. In the absence of tension, this type of translocation can proceed indefinitely; in Figure 6(d) the apparent *≈* 500 nm limitation is due to the length of the DNA being simulated. It has been noted that this does not explain the obstacle bypass events observed in Ref. [10] as a second loop was not observed in Ref. [21].

**Figure 7:**
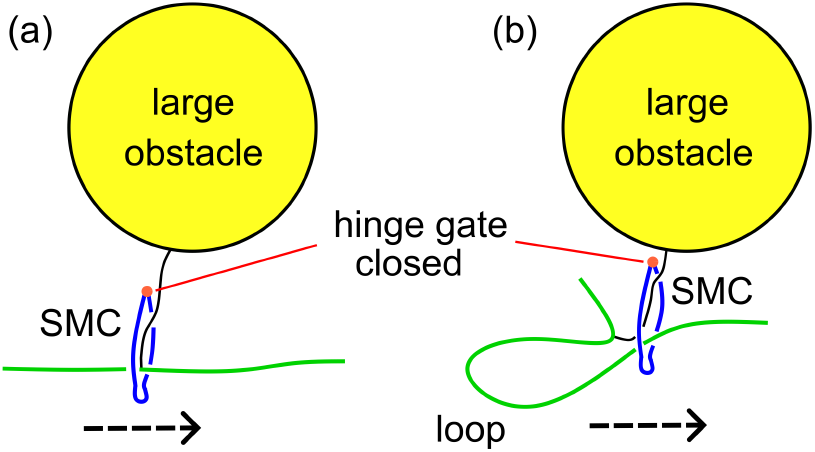
While in some cases the closing of the hinge gate stops the translocation process (a), instances are observed in which the translocation continues by capturing segments of DNA beyond the linker attachment point (b). This leads to the trajectories showing translocation beyond the linker-attachment point of Fig. 6(d).

### 3.3 Obstacle crossing through lumen

We also considered smaller obstacles directly bound to DNA without linkers. For simplicity, we refer to these particles as “nucleosomes”, although they more generally represent DNA-binding complexes, such as RNA polymerases, which are sufficiently small to pass through the SMC ring, Fig. 8(a). We investigated translocation through a regular array of nucleosomes with varying spacings and two initial conditions. In the first condition, translocation starts from a nucleosome-depleted region and ends up in a nucleosome-rich region (Fig. 8(b)). In the second condition, translocation takes place entirely within a nucleosome-rich region (Fig. 8(c)). Actual nucleosomes have a diameter of about 11 nm, whereas each SMC coiled-coil has an approximate length of 50 nm. In our MD model, the nucleosomes are approximated as spherical beads with a diameter of 11 nm, making them effectively larger than real nucleosomes, which are roughly cylindrical with a height of only 5 nm. A representative simulation snapshot of DNA translocation through a nucleosome array is shown in Fig. 8(a). In this configuration, the kleisin is folded, corresponding to state 1 of Fig. 2, which results in the capture of a DNA segment with two nucleosomes.

**Figure 8:**
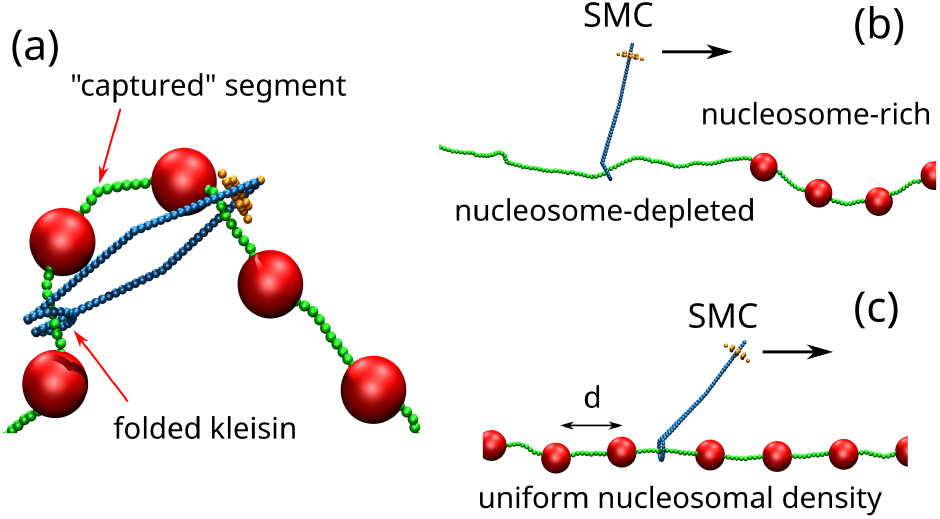
(a) Snapshot of a translocation event with the folding of the kleisin leading to the capture of a DNA segment containing two nucleosomes. The nucleosomes are placed between the DNA strands and bonded to the DNA chain with a bond length proportional to the bead size. The angle potential at the DNA-nucleosome interface is computed in the same way as for the DNA itself, such that the persistence length remains identical. Therefore, there should be no effect of increased stiffness in the simulations. (b,c) Two different initial conditions for the simulations of SMC translocation over DNA with bound nucleosomal-type particles (the arrow indicates the SMC translocation direction). In panel (b) the SMC translocates from a nucleosome-depleted to a nucleosome-rich region. In panel (c) the SMC translocates within a uniform density of nucleosomes.

Figure 9 compares trajectories obtained from three different setups. Panel (a) shows translocation along naked

**Figure 9:**
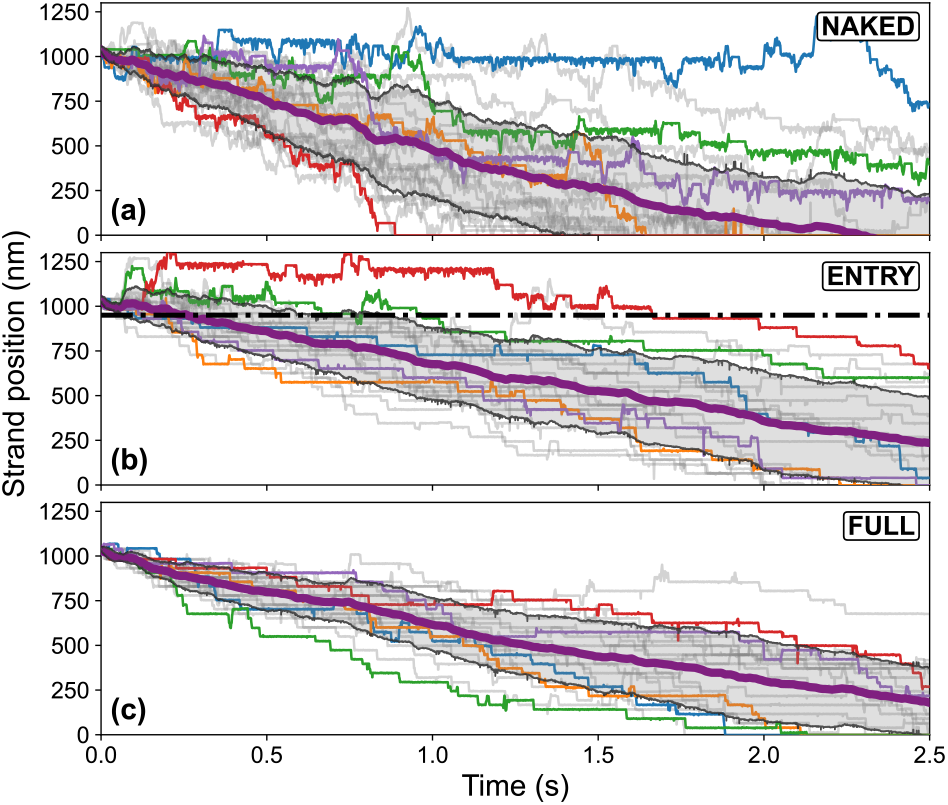
Comparison of simulation trajectories for SMC translocation (a) along naked DNA, (b) from a nucleosome-depleted region to a nucleosome-rich region, and (c) within a region with uniform nucleosomal density. The cases (b) and (c) correspond to the setup shown in Fig. 8(b) and (c), respectively. The thick purple line shows the average SMC position over all trajectories and the gray area represents an interval of *±*1 standard deviation. Note that for (b) and (c), the SMC position is measured in terms of the fiber length, and therefore does not include the contour length of DNA wrapped around nucleosomes.

DNA and serves as a reference, while panels (b) and (c) correspond to DNA containing nucleosomal particles, as in Fig. 8(b) and (c), respectively. This comparison reveals that translocation along naked DNA (a) exhibits the largest variability among individual trajectories. This intrinsic variability arises from fluctuations in the length of DNA segments captured by the SMC complex in state 1. In state 2, the DNA is unbound and temporarily undergoes free diffusion, resulting in a broad distribution of step sizes, including frequent backward steps [14]. In contrast, translocation along nucleosome-decorated DNA (Fig. 9(b,c)) exhibits reduced variability. The nucleosome beads act as obstacles, which limit both the captured segment size during state 1 (hence the lower average translocation speed) and the forward and backward steps during state 2 (hence the narrower step-size distribution). The horizontal dotted-dashed line of Fig. 9(b) indicates the boundary between the nucleosome-depleted and nucleosome-rich region, and reveals that many trajectories initially “bounce off” the interface, performing backward steps before successfully entering the nucleosome-rich region.

Figure 10 shows plots of the mean translocation step size (left axis) and rate (right axis) as a function of the inter-obstacle spacing, *d*, for obstacles with diameters ranging from *D* = 10 nm (i.e., matching that of nucleosomes) up to *D* = 22 nm. The translocation takes place in a uniform density of obstacles, as shown in Fig. 8(c). We note that the translocation rate rapidly drops with decreasing spacing *d* between nucleosomal particles. The translocation nearly comes to a halt below *d ≈* 6 nm for the obstacle of diameter *D* = 18 nm. Larger obstacles with *D ≥* 22 nm are unable to pass through the SMC ring, resulting in a vanishing translocation rate, independent of the interval *d*. The blocking of translocation for small inter-obstacle distance *d* agrees with the experimental result that a dense array of DNA-bound proteins can effectively stop loop extrusion [26].

**Figure 10:**
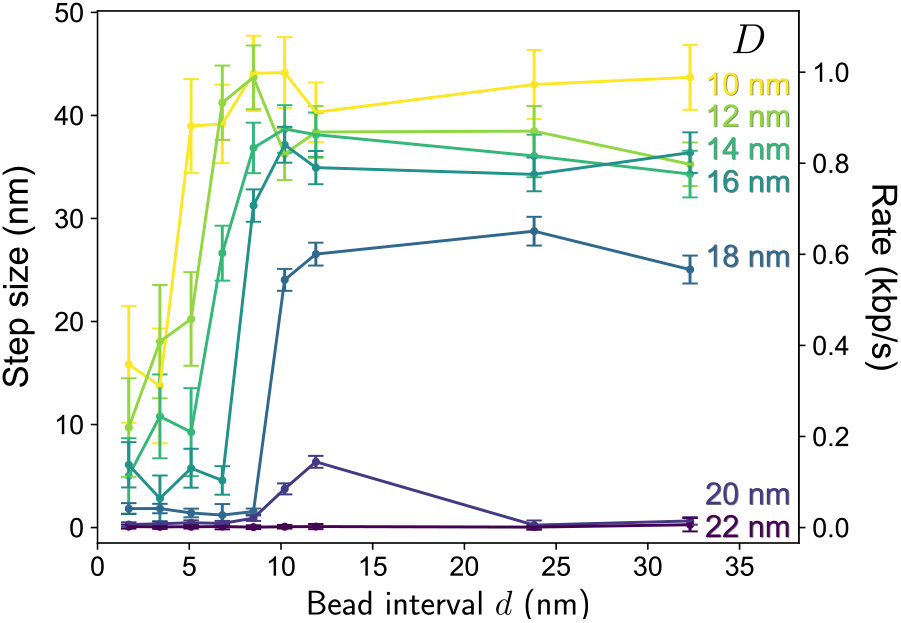
Mean translocation step size (left axis) and corresponding rate (right axis) versus the inter-obstacle interval *d*, as shown in Fig. 8(c), and for varying obstacle diameter *D*. Obstacles with a diameter exceeding 20 nm are no longer able to fit through the SMC ring. As the obstacle spacing decreases below *d ≈* 7 nm the translocation process rapidly stalls, with the drop becoming more abrupt for larger obstacles. The stalling of loop extrusion was experimentally observed for condensin attempting to traverse dense arrays of DNA-bound proteins [26].

### 3.4 Translocation as a function of DNA persistence length

In the model discussed in this work, the SMC complex captures a DNA segment by folding the DNA within its lumen. The SMC arms have a length of approximately 50 nm, which is comparable to the DNA persistence length. Consequently, the captured segment adopts a bent, arc-like conformation [14], which is characteristic of semiflexible polymers that are deformed over contour lengths on the order of their persistence length. We note that DNA used in single-molecule experiments is often visualized using fluorescent dyes as intercalators [8], which may strongly influence DNA mechanical properties. The widely used SYTOX Orange dye has been reported to significantly reduce DNA persistence length from the normally observed value *l*_*p*_ *≈* 50 nm down to *l*_*p*_ *<* 20 nm [27]. To investigate how these factors influence DNA translocation, we carried out a series of simulations spanning a wide range of persistence lengths.

Figure 11 compares the mean step size and corresponding mean rate for DNA polymers of different persistence lengths as a function of *f*, the applied tension at the two ends of the molecules. Runs were performed with forces ranging from *f* = 0 pN to *f* = 4 pN, sampling 20 individual simulations for each force value. The 50 nm DNA mean translocation rate is similar to that reported in Ref. [14] and shows a monotonic decrease of the translocation rate as a function of *f*. As already discussed in Ref. [14], SMC translocation does not stall at high forces due to an “inchworm” mode of translocation (note, however, that LE stalls as discussed in Ref. [14]). The translocation rate is similar for values in the range 10–100 nm. Only at an extremely small persistence length of 1 nm (similar to the value of ssDNA, but admittedly difficult to achieve by chemical modification of dsDNA) does the translocation rate become significantly faster. The smaller persistence length allows a random coil “cloud” of nucleotides to be captured when the kleisin folds.

**Figure 11:**
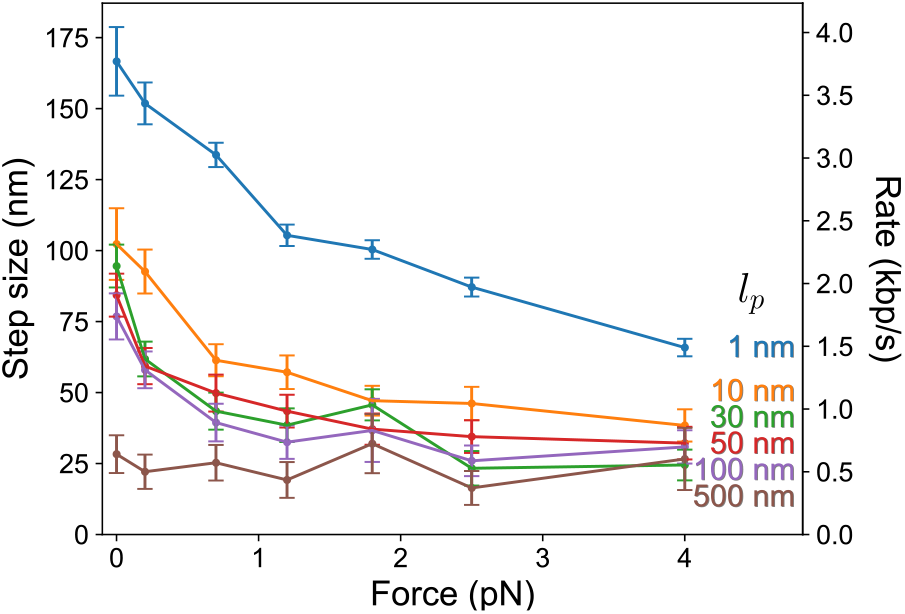
Mean translocation step size (left axis) and corresponding rate (right axis) under different applied forces, shown for values of the persistence length *l*_*p*_ ranging from 1 nm to 500 nm. The given force is applied along the xaxis of the simulation box, simultaneously on the left and right ends of the DNA, such that the net force on the DNA is zero. The reference value of *l*_*p*_ = 50 nm was used for DNA in all other simulations.

## 4 DISCUSSION

A central open question in the current discussion of loop-extrusion mechanisms concerns whether the extruded DNA strands are threaded through the SMC ring itself, consistent with topological and pseudo-topological models (Fig. 1(c,d)), or instead remain external to the ring and are engaged via transient contacts, as posited by nontopological models (Fig. 1(e)). Experimental data provide partial support for both classes of models; hence, the mechanism of SMC-DNA interaction remains debated. On the one hand, there is compelling evidence that DNA can be topologically entrapped by cohesin, condensin, and bacterial SMC complexes, which would favor a (pseudo)topological mechanism of LE. On the other hand, singlemolecule in vitro assays have demonstrated that loop extrusion can proceed even in the presence of steric obstacles larger than the SMC ring diameter [10], a result that was interpreted as supporting a non-topological mode of SMC-DNA interaction.

Recent experiments on bacterial Wadjet, however, demonstrated that large obstacles tethered to an extruding double-stranded DNA (dsDNA) molecule via a thin single-stranded DNA (ssDNA) or RNA linker do not impede loop extrusion, whereas extrusion is stopped when the tether is dsDNA [11]. The hinge site at the apex of the ring was shown to permit passage of thin linker molecules such as ssDNA, but not of thicker ones such as dsDNA [11]. The Wadjet experiments have provided additional support for (pseudo-)topological models of LE. Motivated by these recent findings, we employ molecular dynamics simulations to investigate the properties of the (pseudo-)topological segment-capture model of SMC-mediated LE [14, 17], with particular emphasis on obstacle-crossing and directional-inversion mechanisms, and compare the resulting predictions with the available experimental data. We discuss our findings here and consider alternative nontopological mechanisms that could be consistent with experiments.

### 4.1 Symmetric loop extrusion via strand swapping

The segment-capture model [17] can be adapted to account for both topological and pseudo-topological modes of LE. In the topological case, one dsDNA molecule is threaded through the SMC ring, while the second strand is threaded through another SMC ring in the case of dimeric SMC complexes or is non-covalently tethered to the exterior of the ring via a mechanism commonly referred to in the literature as the “safety belt”. In the pseudo-topological case, by contrast, two dsDNA molecules are simultaneously threaded through the SMC ring. In the topological setup of the segment-capture model the SMC complex functions as a fully asymmetric loop extruder, with one strand remaining permanently anchored and the other being progressively reeled into the SMC ring.

It is known that condensin functions as an asymmetric loop extruder, while cohesin exhibits symmetric LE activity. Recent single-molecule experiments have clarified that this apparent “symmetric” behavior of cohesin arises from frequent reversals of the strand translocation: while each elementary translocation step involves only one strand at a time, the translocating strand alternates over time [24]. Hence the translocation remains intrinsically asymmetric, but the strand exchange generates an overall symmetric extrusion [24]. Here, we have demonstrated that the segment-capture model can carry out this type of directional switching through the strand-swapping mechanism illustrated in Fig. 3. The topological nature of the SMC-DNA coupling enables changes in translocation direction while robustly preserving DNA entrapment within the SMC ring.

Experimental data further indicate that, in addition to strand swapping, unidirectional loop growth is intermittently disrupted by periods of free diffusion and slipping [24]. Consistent with these observations, our simulations produce individual translocation trajectories that are highly stochastic: they are characterized by pronounced noise and frequent backward steps during DNA translocation, as shown in Fig. 6. Although wild-type yeast condensin normally generates DNA loops via asymmetric extrusion, disruption of this behavior occurs when the an-chor subunit protein YCG1 is deleted or when mutations impair the high-affinity DNA-binding region of the safety belt [21]. Under these conditions, condensin LE becomes symmetric [21]. This observation closely parallels our simulation results, in which a transition from asymmetric to symmetric LE is induced by weakening the anchor-binding site, thereby increasing the likelihood of dissociation events leading to strand swapping.

A directional strand-swapping mechanism within a nontopological framework, in which the DNA remains external to the SMC ring, entails several conceptual and mechanistic difficulties. In the non-topological model proposed in Ref. [12], the authors postulate that the kleisin subunit encircles the DNA, mediated by the binding of HEAT protein complexes, without the DNA being threaded through the SMC ring. The corresponding architecture is depicted schematically in Fig. 12. HEAT proteins constitute auxiliary regulatory cofactors that promote DNA loading, unloading, translocation, and stable association. These proteins exhibit complex-specific variations and are designated by different names in distinct SMC complexes, for example NIPBL/Scc2/Pds5 in cohesin and Ycg1/Ycs4 in condensin.

**Figure 12:**
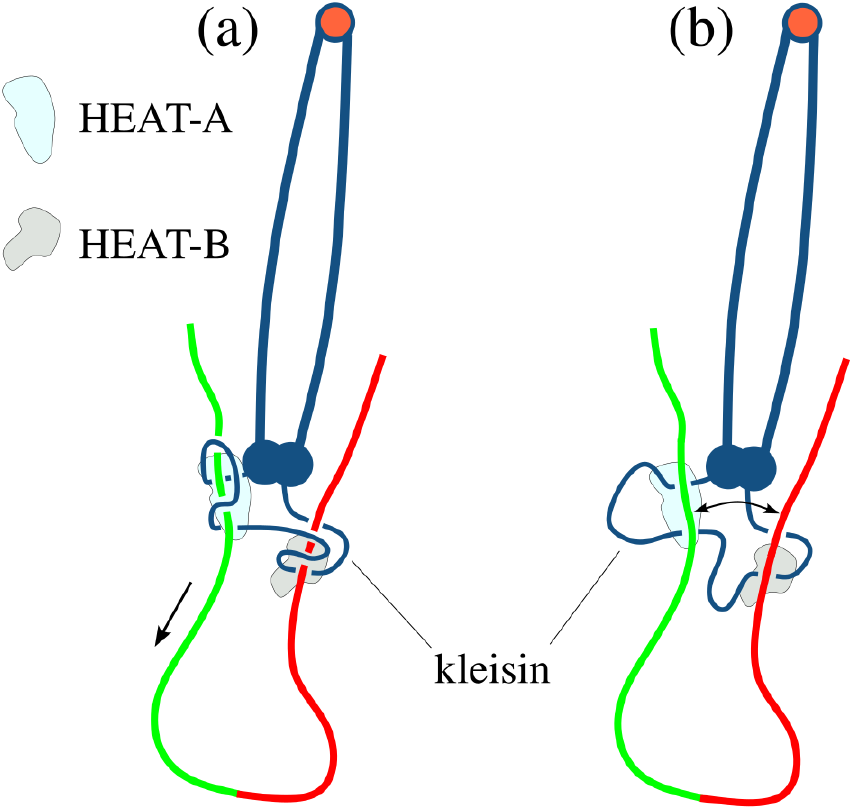
(a) In the non-topological loop-extrusion model described in Ref. [12], the kleisin subunit encircles the DNA with the assistance of regulatory protein complexes denoted HEAT-A and HEAT-B, but the DNA is not threaded through the kleisin. HEAT-A proteins bind in proximity to the SMC head domains and contribute to the translocation of DNA strands through conformational rearrangements (for details of these conformational changes leading to translocation, see Ref. [12]). By contrast, HEAT-B binds stably to DNA and does not directly participate in the extrusion process. Under these conditions, the green DNA strand elongates. (b) To achieve symmetric loop extrusion via growth of the red strand, an additional swapping mechanism is required. If the DNA is not threaded through the SMC complex during this process, the loop can be lost.

Two functional classes of HEAT proteins are typically distinguished. HEAT-A proteins bind predominantly in proximity to the ATPase head domains and participate directly in DNA capture and translocation. In contrast, HEAT-B proteins associate more stably with the kleisin subunit and primarily regulate DNA retention and release. Within the framework of the model of Ref. [12], a strand-swapping process would require an exchange between two DNA segments such that the strand initially bound to HEAT-A (HEAT-B) relocates and re-associates with HEAT-B (HEAT-A), respectively. Although such strand exchange is readily accommodated when the DNA is topologically entrapped by, and threaded through, the SMC ring, it becomes problematic in a non-topological configuration. In the latter case, any transient dissociation of either DNA segment from the kleisin would lead to the loss of the DNA loop, as illustrated in Fig. 12(b). Thus, the proposed strand-switching mechanism appears intrinsically fragile and is unlikely to be robust in a non-topological arrangement.

We note that an alternative explanation for the emergence of symmetric versus asymmetric LE has recently been proposed [28]. In that work, symmetric extrusion is attributed mainly to dimers of SMC complexes, in which two mechanically coupled SMC rings each entrap one DNA arm. Symmetric loop growth then arises from the coordinated action of the two motor complexes, each translocating along a different DNA arm. In contrast, the present work focuses on the behavior of a single SMC complex. It will therefore be interesting to extend the segment-capture model to investigate how mechanical coupling between two SMC complexes influences the dynamics and symmetry of loop extrusion. This may provide microscopic insight into the complex phenomenology observed in [28].

### 4.2 Obstacle crossing via hinge gate

Motivated by experiments investigating LE in the presence of DNA-bound “obstacles”, we performed a series of simulations of DNA translocation with two distinct classes of obstacles. These simulations focus on characterizing how such obstacles affect the dynamics of DNA translocation.

The first class consists of large nanoparticles tethered to the DNA via a thin linker molecule, such as ssDNA, with the simulation setup schematically illustrated in Fig. 5(a). For this class of obstacles, we model translocation through a hinge-gate passage located at the apex of the SMC complex, a mechanism recently identified in studies of the bacterial Wadjet system [11]. Our simulations demonstrate that the thin linker can traverse the hinge gate at a rate that depends sensitively on the gate width. For narrower gate openings, we observe a reduction in the average SMC translocation velocity as the complex approaches the obstacle, followed by an increase in velocity once the obstacle has been passed. This leads to a pronounced curvature in the mean position-time traces shown in Fig. 6(c), which could be tested in single-molecule experiments. Interestingly, we found that for a subset of trajectories translocation does not stop, even when the hinge gate is nominally sealed. This bypass mechanism, illustrated in Fig. 7, is consistent with and provides further support for a scenario previously hypothesized in the literature [14].

### 4.3 Obstacle crossing through SMC lumen

The second class of obstacles investigated consists of particles directly bound to DNA that are sufficiently small to pass through the SMC lumen. Arrays of these “nucleosome-like” particles cause a reduction in the translocation velocity. Our analysis shows, however, that this deceleration is relatively modest for particles with a size comparable to physiological nucleosomes, characterized by a diameter of *D ≈* 10 nm, whereas the impact becomes substantial for larger particles (Fig. 10). When the SMC complex translocates from a region depleted of nucleosomes toward a nucleosome-enriched region, a subset of SMC trajectories is reflected, indicating that a dense nucleosomal array acts as a translocation barrier. If the obstacle density is increased such that the spacing between obstacles becomes smaller than approximately 7 nm (a value that depends on the obstacle size), translocation stops. This is in line with the results of recent experiments on LE by condensin on arrays of telomeric DNA-binding proteins Rap1 [26], which show that dense arrays of Rap1 can completely halt the LE process.

### 4.4 Noise reduction by arrays of obstacles

Another noteworthy observation is that translocation through a chromatin region with a uniform nucleosome density exhibits substantially smaller trajectory-to-trajectory variability than translocation along naked DNA. Segment-capture model simulations of bare DNA [14] generally predict a broad distribution of step sizes, including forward steps of up to 200 nm, far exceeding the SMC ring size (*∼* 50 nm), as well as a significant fraction of backward steps. This broad distribution originates from diffusive motion during the transition from SMC state 2 to state 0, when the captured DNA segment is transferred from the upper compartment to the kleisin. During this stage, the DNA is temporarily unbound from the SMC complex and diffuses freely, effectively smearing out the captured segment length and giving rise to a wide range of translocation step sizes. In the presence of nucleosome-like particles, however, this diffusive motion is strongly constrained by the presence of obstacles that act as transient barriers to free diffusion. As a result, both the spread of step sizes and the variability between individual trajectories are markedly reduced, leading to a more uniform translocation process. It would be interesting to test this prediction in single-molecule experiments.

## 5 CONCLUSION

We have shown that, in addition to naturally explaining symmetric LE via the strand-swapping mechanism, the segment-capture model yields a number of experimentally testable predictions regarding the interaction of SMC complexes with DNA-bound obstacles. For large obstacles attached to the DNA via a flexible linker, the model predicts a transient slowdown upon obstacle encounter, followed by an acceleration of translocation once the obstacle has been bypassed. For nucleosome-like particles that can thread through the SMC ring, the model predicts a pronounced suppression of both step-size fluctuations and trajectory-to-trajectory variability. These effects provide experimental signatures that can be directly probed in single-molecule assays, adding substantially to our understanding of the molecular mechanism of loop extrusion.

## Supporting information

Movie: Asymmetric extrusion

Movie: Symmetric extrusion via strand swapping

## 6 DATA AVAILABILITY

The MD model simulation code is available at https://github.com/LucasDooms/SMC_LAMMPS.

The numerical data and analysis scripts used to produce the figures are available at https://doi.org/10.5281/zenodo.21098585.

## ACKNOWLEDGMENTS

LD acknowledges funding from FWO project G0A8L24N INSITE. JFM acknowledges the support of the US NIH through grant R35-GM161403. SG acknowledges support by the SNSF grant 3200-0-239885.

