## Supplemental for "Obstacle bypassing and intrinsically asymmetric loop extrusion in the segment-capture model of Structural Maintenance of Chromosomes complexes"

### S1 Movies

We include two movies of molecular dynamics simulations illustrating loop extrusion by the segment-capture model. Movie S1 shows asymmetric loop extrusion, in which one of the two DNA strands remains permanently attached to the SMC binding site. Movie S2 shows a simulation in which this DNA strand undergoes periodic detachment from the binding site. In the former case the loop grows asymmetrically, whereas in the latter the repeated attachment and detachment events lead to effectively symmetric loop extrusion. Figures S1 and S2 show the initial and final configurations of the two simulations. As in the main text, we divided the DNA strand of the loop in two colors. In the asymmetric case of Fig. S1 only the red strands grows.

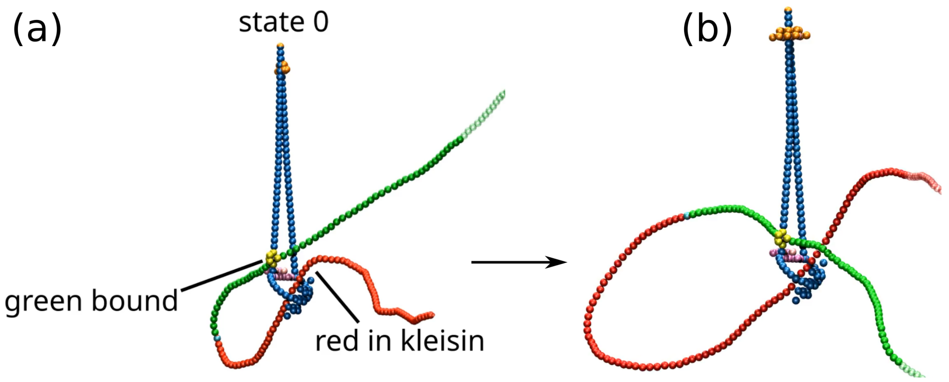


Figure S1: Movie S1 shows a molecular dynamics simulation of the segment-capture model with (a) the initial configuration and (b) the final configuration. During the simulation, the green DNA strand remains permanently bound to the SMC complex. Folding of the kleisin subunit captures the red DNA strand, resulting in asymmetric loop extrusion where only the red strands grow.

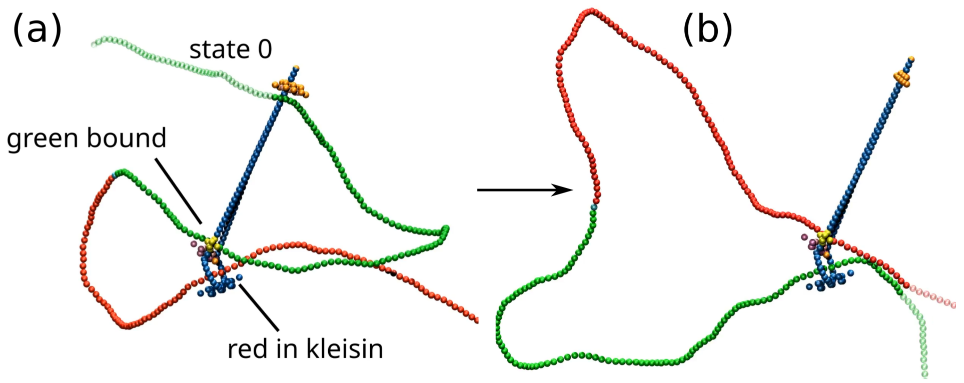


Figure S2: Movie S2 shows a molecular dynamics simulation of the segment-capture model in which the bound DNA strand periodically detaches from the SMC complex. Panels (a) and (b) show the initial and final configurations, respectively. While in (a) the green strand is bound, in (b) the red one is bound and the green one in kleisin. Although each individual translocation step is intrinsically asymmetric, the repeated detachment events give rise to effectively symmetric loop extrusion.
